# qpMerge: Merging different peptide isoforms using a motif centric strategy

**DOI:** 10.1101/047100

**Authors:** Matthew M. Hindle, Thierry Le Bihan, Johanna Krahmer, Sarah F. Martin, Zeenat B. Noordally, T. Ian Simpson, Andrew J. Millar

**Author notes:** Corresponding author: Andrew J. Millar, SynthSys and School of Biological Sciences, University of Edinburgh, UK. EH9 3JD. Abbreviations:* AC: Alternative Cleavage site, AM: Alternative Modifications, CS: Charge State, CV: Coefficient of Variation, EMM: Exact Motif Matches, SD: Standard Deviation.

## Abstract

Accurate quantification and enumeration of peptide motifs is hampered by redundancy in peptide identification. A single phosphorylation motif may be split across charge states, alternative modifications (*e.g.* acetylation and oxidation), and multiple miss-cleavage sites which render the biological interpretation of MS data a challenge. In addition motif redundancy can affect quantitative and statistical analysis and prevent a realistic comparison of peptide numbers between datasets. In this study, we present a merging tool set developed for the Galaxy workflow environment to achieve a non-redundant set of quantifications for phospho-motifs. We present a Galaxy workflow to merge three exemplar dataset, and observe reduced phospho-motif redundancy and decreased replicate variation. The qpMerge tools provide a straightforward and reusable approach to facilitating phospho-motif analysis.

The source-code and wiki documentation is publically available at http://sourceforge.net/projects/ppmerge. The galaxy pipeline used in the exemplar analysis can be found at http://www.myexperiment.org/workflows/4186.

## Introduction

Accurate quantification of protein motifs that are post-translationally modified is one of the most common proteomics strategies. However, motif redundancy among detected peptides by mass spectrometry (MS) leads to errors and ambiguities in the analysis. The detected quantification of some motifs can be fragmented across multiple MS features. The representation of phospho-motifs by several variants can result from: (1) different charge states, (2) alternative modifications (oxidation and acetylation to name a few), and (3) multiple sites of protease digestion. When a single motif is redundantly represented across several peptides it can affect the sample statistics through fragmentation of observations (*e.g.* ANOVA, *t*-tests), and population statistics by the distortion of the overall distribution (*e.g. z*-ratios, GO enrichment). It also hinders the comparison across datasets, as the quantity and quantification of motifs are biased by variable fragmentation, which is highly dependent on the biological sample and experimental protocol. Finally, this information redundancy is confusing for any biological interpretation.

Many of the published analyses of quantitative proteomics datasets use complete software packages like Progenesis (Nonlinear Dynamics), PEAKS [1] or MaxQuant [2] to name few. However, there is also a growing trend for the more configurable and extensible workflow frameworks such as the Trans-proteomics pipeline (TPP), OpenMS [3], and GalaxyP. The Galaxy [4] framework has a wide user base in the genomics community and allows workflow composition using a large tool-shed of existing bioinformatics tools. Workflows can be shared and re-used through MyExperiment [5]. The msCompare tool has previously demonstrated quantitative-proteomics analysis within Galaxy [6]. Data standards are vitally important to promote data and tool re-use. Standards for quantitative proteomics data formats are still in the early adoption phase and formats like MzQuantML [7] are not yet widely supported.

One approach to achieving a non-redundant quantitation of phospho-motifs is to quantify each peptide but only assign a reference peptide, according to some quality criteria, as the quantification of the motif. This discards peptides, redundant to the phospho-motif reference, which could have a significant contribution to the quantification of the motif. Alternatively, Casado *et al.* [8] quantified the abundance of phospho-motifs as the mean quantification across all matching peptides. In most quantitative proteomics analyses, protein abundance is predominantly defined as the intensity sum of all peptides comprising the protein; in the qpMerge approach we extend this concept to the quantification of phospho-motifs. Using the sum avoids the disproportionate effect of low abundance peptides on the mean.

## Methods

We implemented the qpMerge set of tools as an open source Java library that can be used programmatically, as a command line tool or on a Galaxy server by installing the provided tool wrappers. Supporting workflow composition in Galaxy promotes component re-use, reproducible analysis, and importantly provenance tracking. The workflow for generating the exemplar analysis in this paper (implemented in Galaxy) is publically shared in MyExperiment. Fig. 1A outlines the core processing steps to remove redundant phospho-motifs. Each step is wrapped as a plugin which is enacted sequentially within in a Galaxy workflow. We augmented this core workflow with tools for data import (MzQuantML, Progenesis and Maxquant), export (xml, tabular and MzQuantML), and annotation (domains, accessions, and statistics). The pipeline also incorporates components for statistical analysis, domain annotation through InterProScan 5 [9], and accession mapping through the UniProt and NCBI web services. Our implantation is in Java and implements a generic and reusable data-model that tracks the provenance of a phospho-motif back to the quantification of individual peptides. Merged phospho-motifs are represented as a graph, retaining the incorporated peptides as parent nodes. Tracking the provenance of a given phospho-motif entails traversing up the hierarchy of peptides that was constructed by successive merging steps.

**Figure 1.**
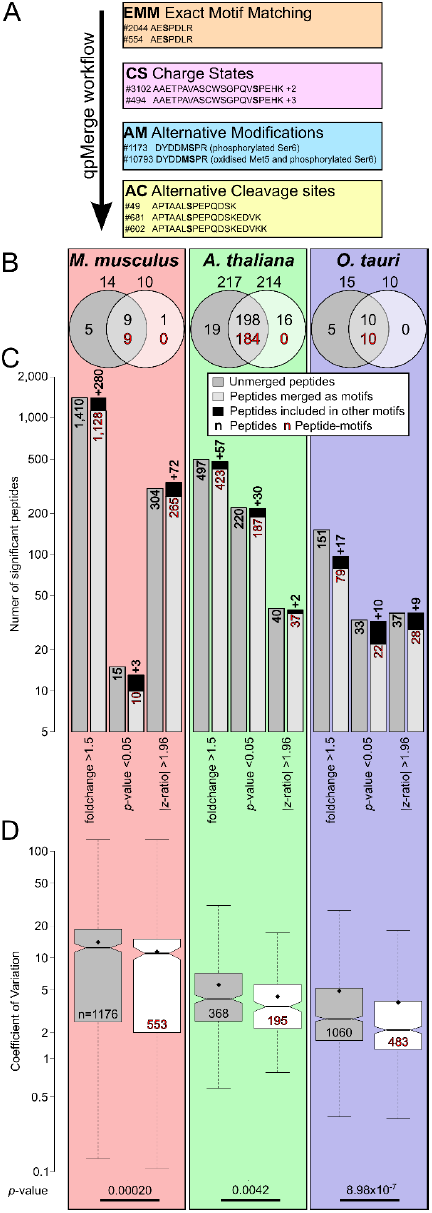
Impact of the qpMerge pipeline on statistical analysis. (A) A summary schematic of the core pipeline with examples of peptides that are merged from the *Mus musculus* dataset. (B) Venn diagrams for each experiment describing the number and intersection of peptides and motifs that have statistically significant differences in mean quantification and fold-change > 1.5, prior to and after the merging process. The number of phospho-motifs after merging is given in brackets. (C) Peptides that are above the individual thresholds for statistical significance (*t*-test *p*-value of < 0.05; a magnitude of fold change >1.5; |z-score| > 1.96). (D) The overall effect of the merging process on the distribution of the CV across the three datasets. The CV is shown for all replicates in both control and treatment conditions. Non-overlapping notches indicate statistically significant differences in the black median line. Peptide quantifications affected by the respective merging step, and included in the boxplots, are indicated by the number in the lower quartile. The statistical significance of the difference of the mean as indicated by a Mann-Whitney U test on the log normalised CV.

We provide an exemplar workflow in Galaxy that demonstrates the merging of fragmented phospho-motifs based on a conjoint Progenesis and Maxquant inputs. We report the impact of the pipeline on the analysis of three published datasets from *Mus musculus* [10], *Arabidopsis thaliana* [11] and *Ostreococcus tauri* [12].

In our three examples we import quantification from Progenesis and peptide modification predictions (together with their probabilities) from MaxQuant. Parameters for these tools have previously been described [13]. For each dataset the workflow requires metadata on the experiment, variables, authors, and observations. This can be provided in a flat-file, XML-file or Galaxy form. The initial import component entails (1) integration of the two datasets and (2) removing redundancy within individual peptide identifications. MaxQuant phospho-peptide assignments are mapped onto Progenesis quantitation based on sequence and mass. Where multiple sequences are listed for a single identification, we retain peptides within 10% of the best Mascot score.

As part of the qpMerge process, we define a phospho-motif as the set of phosphorylated amino acids that are co-observed (appear on a single peptide) on a protein. We aim to eliminate or combine all peptides that result in redundant occurrences of a phospho-motif. Peptides that have different sites or quantities of phosphorylated amino acids are considered distinct. The first component of the workflow is filtering Exact Motif Matches (EMM). These are peptides with the same sequence, modification, and charge. The new phospho-motif retains the quantification of the highest scoring peptide. The addition of MaxQuant phospho-site predictions allows the grouping of peptides with a similar mass that are assigned distinct modification sites in Progenesis but have non-differentiable probabilities for residues sites in MaxQuant. The second stage in the workflow is merging the different Charge States (CS). These are peptides that have the same sequence and modification but a different charge. The new phospho-motif sums the intensity of the merged charge-variants. The parameters of the Alternative Modifications (AM) allow the merging of modification that are not part of the motif of interest. In our exemplar workflow we group peptides with the same phospho-motifs but differing oxidation and acetylation states. We sum the quantification across peptides grouped by AM. Alternative Cleavage sites (AC) are produced when a given protease produces strand breaks across multiple sites in a protein which has the consequence a given specific motif found in several peptides of different length. We aligned peptides with unique phospho-motifs, allowing for overlap at N-and C-termini. The aligned peptides quantifications are summed into the new phospho-motif.

All three published exemplar datasets represent pairwise comparisons of a treatment variable against a control. They were selected because they represent diverse eukaryotic taxa (Plantae, Algae and Metazoa), and experimental conditions. The *A. thaliana* dataset compares two detergents for use in protein extraction: sodium dodecyl sulfate and octyl glucoside. The *O. tauri* dataset contrasts algae cells with and without the application of the kinase inhibitor IC-261. The *M. musculus* dataset examines the FD6 cell-line before and after the application of Fibroblast Growth Factor 4 (FGF4). All datasets were mean normalized and archsinh transformed. We applied homoscedastic *t*-tests for *A. thaliana* and *O. tauri*, and a paired *t*-test for *M. musculus*. The *z*-ratios were calculated, as described by Cheadle *et al.* [14]. Magnitude of fold-change is the ratio of the largest to the smallest sample-average in the non-transformed data.

## Results

In all three of the exemplar experiments the process of merging reduced the final set of phospho-motifs. Within the *M. musculus* experiment 1,595 phospho-peptides were reduced to 1,267 phospho-motifs, similarly *A. thaliana* was reduced from 735 to 645, and *O. tauri* from 1,312 to 1,004. There were also changes to phospho-motifs with statistically significant changes in quantification (combined threshold statistic; *p*-value < 0.05 and fold change > 1.5) (Fig. 1B). For *A. thaliana* this resulted in a reduced number of reported motifs that summarised 198 significant peptides into 184 phospho-motifs. In *M. musculus*, *A. thaliana* and *O. tauri* datasets the merging process excluded 5, 19, and 5 peptides respectively. The workflow did not result in the inclusion of previously insignificant phospho-motifs; however the merged set of significant phospho-motifs incorporated quantification from an additional *M. musculus* peptide and 16 *A. thaliana* peptides. While the overall phospho-motif redundancy is reduced the peptides supporting these significant phospho-motifs are generally similar before and after merging. We broadened the analysis to examine the impact of merging on individual statistical tests (*t*-test, fold-change, *z*-ratio). As with the combined statistic, in all statistical tests and experiments the number of phospho-motifs was reduced after merging (Fig. 1C). The number of underlying peptides supporting the significant phospho-motifs were also consistently smaller with the exception of the *z*-ratio for *M. musculus* where 304 significant peptides were reduced to 265 phospho-motifs that represented 337 peptides. The largest fall in peptide significance was fold-change in *O. tauri*, where merging reduced 151 peptides to 79 phospho-motifs (representing 96 peptides). In each of the three datasets we also observed a reduction in the mean coefficient of variation (CV) within the merged phospho-peptides compared to the unmerged peptides (Fig. 1D). CV differences between the phospho-motifs and their corresponding pre-merged peptides also consistently show a reduction in CV upon merging. *M. musculus* has a mean CV difference of −2.53 (SD, 7.53), *A. thaliana* −1.13 (SD, 2.02) and *O. tauri* −0.91 (SD, 3.11). This indicates that the merging process reduces replicate variation with respect to phospho-motifs.

In order to dissect out the impact of each step in the workflow we individually applied each component of the workflow to the datasets. We found that AM impacted the most peptides, with 928 peptides affected in *M. musculus* (Fig. 2A), 284 in *A. thaliana* (Fig. 2B) and 796 in *O. tauri* (Fig. 2C). AC and CS had the next greatest impact on peptide numbers and EMM the least. With one exception (EMM in *A. thaliana*), all steps resulted in a reduction in CV. AM and AC showed statistically significant (*p*-value < 0.05) reductions in CV in all three experiments.

**Figure 2.**
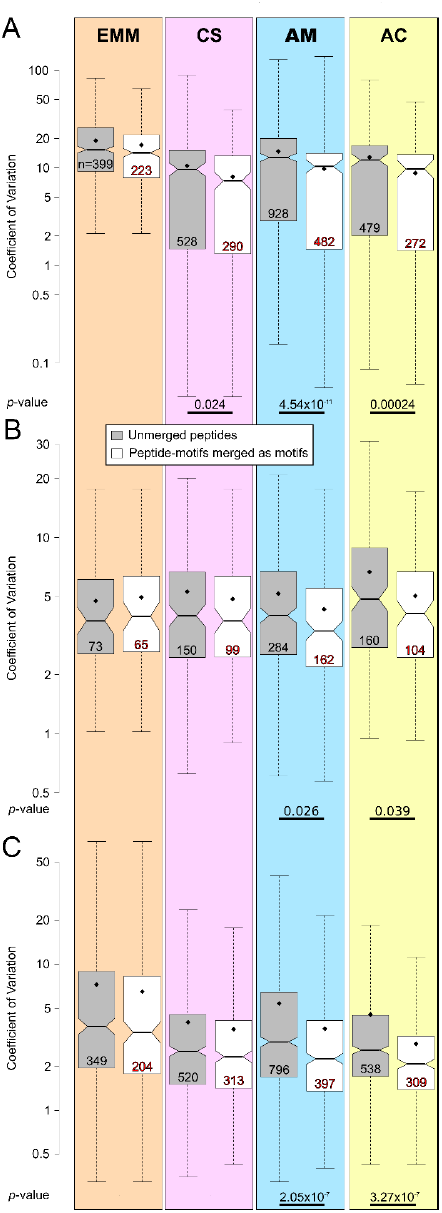
Contribution of each step in the qpMerge pipeline to final phospho-motif counts and CV. The effect on the CV of merged peptides for each step in the pipeline applied independently for the (A) *M. Musculus* (B) *A. thaliana* (C) *O. tauri*. The black dot represents the mean. In the lower quartile of each boxplot the number of peptides affected by the merging process is given.

## Conclusions

In summary, the qpMerge tool provides a set of post-processing algorithms that reduces the phospho-motif redundancy within a dataset. Reduced redundancy facilitates further analyses with respect to phospho-motifs (*e.g.* logo identification and enrichment [15]) and provides a more consistent measure with which to compare datasets. This merging process can affect common measures of statistical and biological significance, and we observed an overall trend to reduce replicate variation within three diverse exemplar datasets.

## Acknowledgements

Funding: Supported by BBSRC and EPSRC awards BB/D019621 and BB/J009423.

## References

[1] Zhang, J., Xin, L., Shan, B., Chen, W., et al., PEAKS DB: de novo sequencing assisted database search for sensitive and accurate peptide identification. Mol. Cell. Proteomics MCP 2012, 11, M111.010587.

[2] Cox, J., Mann, M., MaxQuant enables high peptide identification rates, individualized p.p.b.-range mass accuracies and proteome-wide protein quantification. Nat. Biotechnol. 2008, 26, 1367–1372.

[3] Weisser, H., Nahnsen, S., Grossmann, J., Nilse, L., et al., An automated pipeline for high-throughput label-free quantitative proteomics. J. Proteome Res. 2013, 12, 1628–1644.

[4] Goecks, J., Nekrutenko, A., Taylor, J., Galaxy: a comprehensive approach for supporting accessible, reproducible, and transparent computational research in the life sciences. Genome Biol. 2010, 11, R86.

[5] Goble, C.A., Bhagat, J., Aleksejevs, S., Cruickshank, D., et al., myExperiment: a repository and social network for the sharing of bioinformatics workflows. Nucleic Acids Res. 2010, 38, W677–682.

[6] Hoekman, B., Breitling, R., Suits, F., Bischoff, R., Horvatovich, P., msCompare: a framework for quantitative analysis of label-free LC-MS data for comparative candidate biomarker studies. Mol. Cell. Proteomics MCP 2012, 11, M111.015974.

[7] Walzer, M., Qi, D., Mayer, G., Uszkoreit, J., et al., The mzQuantML data standard for mass spectrometry-based quantitative studies in proteomics. Mol. Cell. Proteomics MCP 2013, 12, 2332–2340.

[8] Casado, P., Alcolea, M.P., Iorio, F., Rodríguez-Prados, J.-C., et al., Phosphoproteomics data classify hematological cancer cell lines according to tumor type and sensitivity to kinase inhibitors. Genome Biol. 2013, 14, R37.

[9] Jones, P., Binns, D., Chang, H.-Y., Fraser, M., et al., InterProScan 5: genome-scale protein function classification. Bioinformatics 2014, btu031.

[10] Docherty, F.M., The identification of FGF-dependent phosphorylation events in embryonic stem cells using mass spectrometry. Biosci. Horiz. 2010, 3, 21–28.

[11] Krahmer, J., Hindle, M.M., Martin, S.F., Le Bihan, T., Millar, A.J., in:, Sehgal A (Ed.), Academic Press, Circadian Rhythms and Biological Clocks 2014.

[12] Van Ooijen, G., Hindle, M., Martin, S.F., Barrios-Llerena, M., et al., Functional Analysis of Casein Kinase 1 in a Minimal Circadian System. PLOS ONE 2013, 8, e70021.

[13] Van Ooijen, G., Martin, S.F., Barrios-Llerena, M.E., Hindle, M., et al., Functional analysis of the rodent CK1tau mutation in the circadian clock of a marine unicellular alga. BMC Cell Biol. 2013, 14, 46.

[14] Cheadle, C., Vawter, M.P., Freed, W.J., Becker, K.G., Analysis of Microarray Data Using Z Score Transformation. J. Mol. Diagn. JMD 2003, 5, 73–81.

[15] O’Shea, J.P., Chou, M.F., Quader, S.A., Ryan, J.K., et al., pLogo: a probabilistic approach to visualizing sequence motifs. Nat. Methods 2013, 10, 1211–1212.

